# RSAT Var-tools: an accessible and flexible framework to predict the impact of regulatory variants on transcription factor binding

**DOI:** 10.1101/623090

**Authors:** Walter Santana-Garcia, Maria Rocha-Acevedo, Lucia Ramirez-Navarro, Yvon Mbouamboua, Denis Thieffry, Morgane Thomas-Chollier, Bruno Contreras-Moreira, Jacques van Helden, Alejandra Medina-Rivera

**Author notes:** To whom correspondence should be addressed., Correspondence may also be addressed to Jacques van Helden.

## Abstract

Gene regulatory regions contain short and degenerated DNA sites recognized by transcription factors (TFs). When such regions harbor SNPs, the DNA motifs where TFs bind may be affected, thereby altering the transcriptional regulation of the target genes. Such regulatory SNPs have been implicated as causal variants in GWAS studies. In this study, we describe the application of the programs *Var-tools* designed to predict regulatory variants, and present four case studies to illustrate their usage and applications. In brief, *Var-tools* facilitate i) obtaining variation information, ii) interconversion of variation file formats, iii) retrieval of sequences surrounding variants, and iv) calculating the change on predicted TF affinity scores between alleles, using motif scanning approaches. Notably, the tools support the analysis of haplotypes. The tools are included within the well-maintained suite Regulatory Sequence Analysis Tools (RSAT, http://rsat.eu), and accessible through a web interface that currently enables analysis of five metazoa and ten plant genomes. *Vart-tools* can also be used in command-line with any locally-installed Ensembl genome. Users can input personal collections of variants and motifs, providing flexibility in the analysis.

## Introduction

Non-coding DNA sequence harbors the gene regulatory information necessary for spatial and temporal gene expression patterns. Specifically, gene regulatory regions are short, highly redundant DNA motifs recognized by transcription factors (TFs). These regulatory regions may contain genetic variants, Single Nucleotide Polymorphisms (SNPs) or indels, that alter the DNA TF binding site (TFBS), and thereby the binding of TFs. Moreover, it has been reported that 93.7% of variants that have been associated to human traits or diseases have been found to be located in non-coding regions [1,2], and particularly enriched in open chromatin regions [3], indicating that these variants may have an effect on regulatory mechanisms, which could explain the observed phenotypes.

The Regulatory Sequence Analysis Tools (RSAT, http://rsat.eu) [4,5] has established itself in the last 20 years as a major software suite dedicated to the analysis of regulatory regions, with five public servers supporting more than 500 eukaryote and 9000 prokaryote genomes. With a major focus on usability and accessibility to users with and without formal bioinformatics training, RSAT provides tools to retrieve sequences, perform motifs analysis, evaluate TF motif quality, compare and cluster motifs, convert file formats, etc. Here we describe *Var-tools*, a subset of tools included in RSAT that enable users to analyse regulatory variants and assess their putative impact on TF binding sites.

### Current approaches for detecting potential regulatory variants

The fact that many variants are located in non-coding regions triggered the development of bioinformatic tools focused on identifying the regulatory potential of these genetic variants. Starting from a list of SNPs, bioinformatics analyses can help formulating hypotheses on which TFs may be impacted by a genetic variant. However, there are numerous challenges for *in silico* analysis to unravel the impact of genetic variations in gene regulatory regions. Several tools and resources have been published, providing alternative methods to tackle this problem (Table 1). Most of them are either based on pattern-matching approaches to evaluate the impact of alleles on TF binding using changes on scores, or on machine learning models built using functional annotations of the regulatory regions, *e*.*g*. epigenomics and transcriptomics data. Still, these resources and tools have limitations hampering their usage in several organisms [6–8], on new annotated variants [9–11], and/or on analyses with personal collections of TF motifs [6,7,12].

A first set of tools (labeled ML in Table1) aim for the identification of potential regulatory variants by integrating several types of data, beyond taking into account potential disruption of TF binding. Particularly, [13] integrated DNA-seq data with SVM approaches to identify variants that could potentially disrupt TF binding.

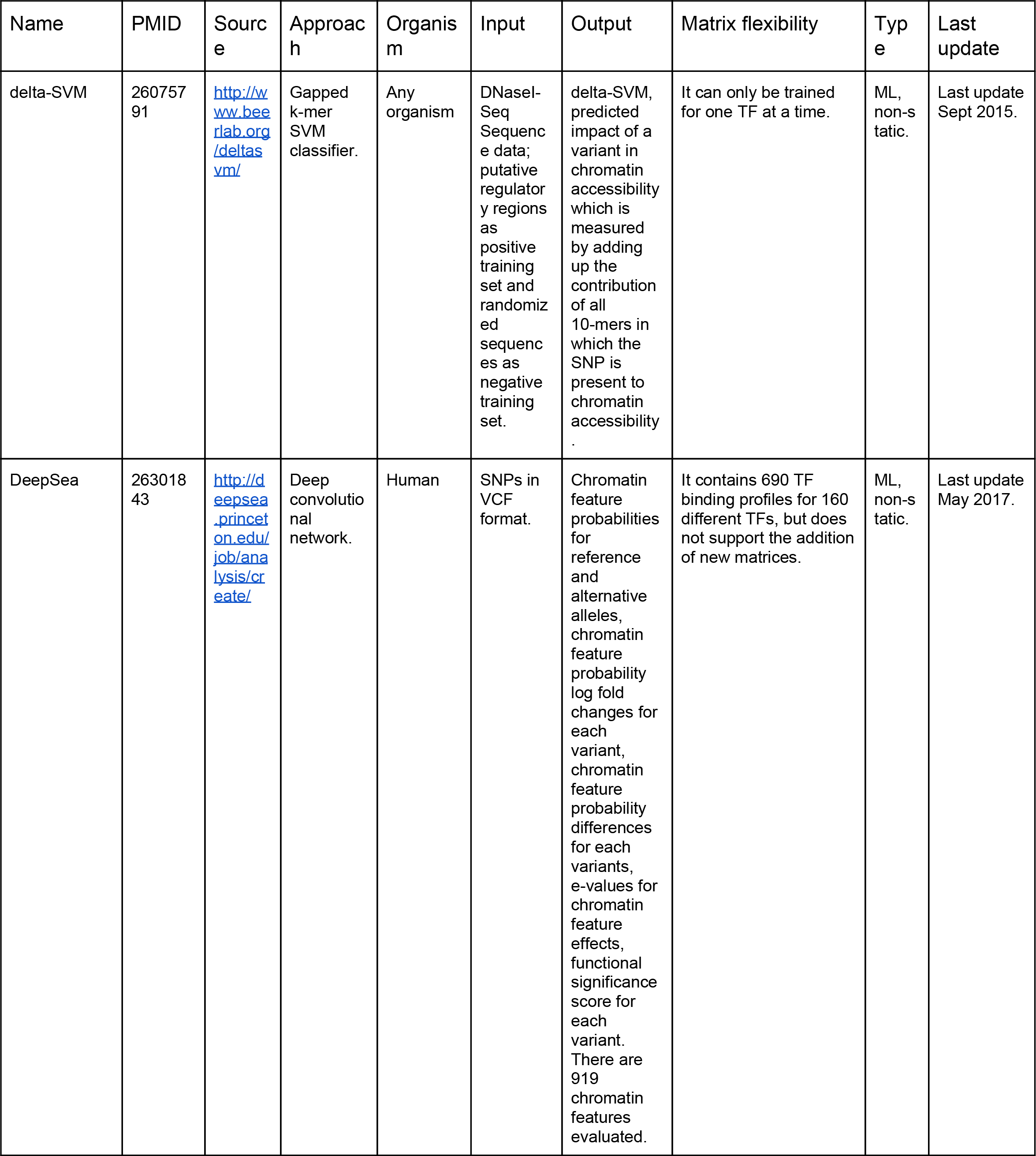
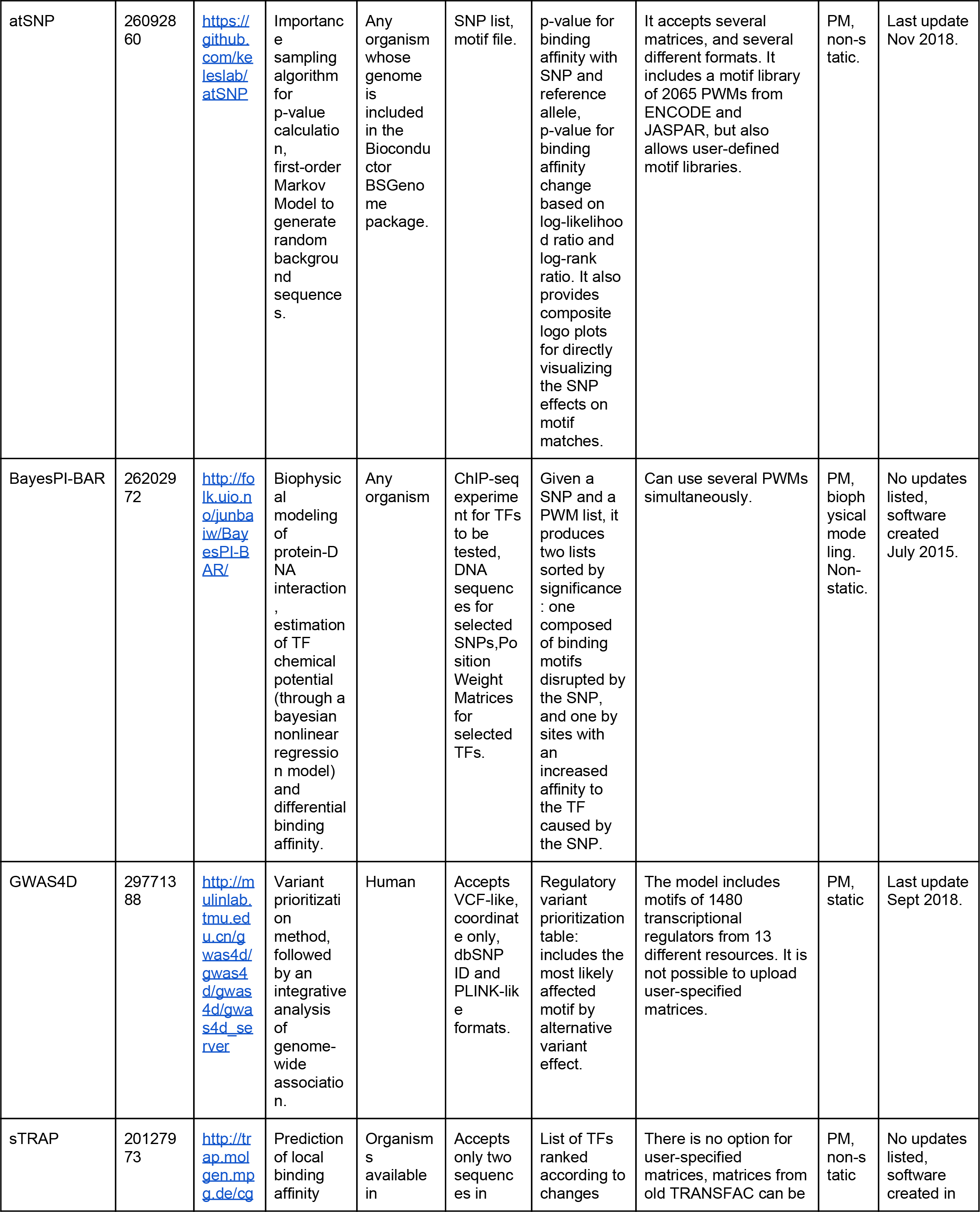
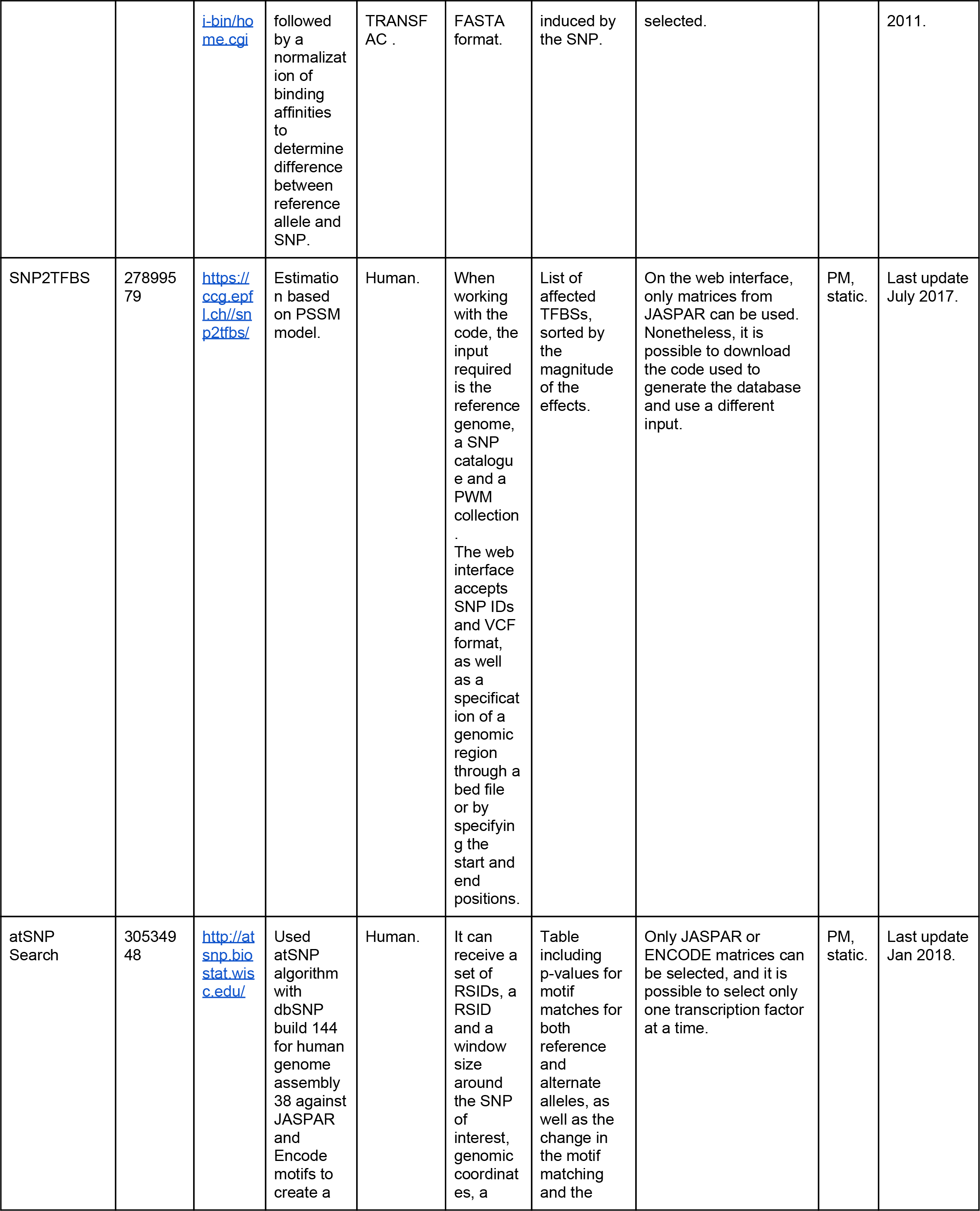
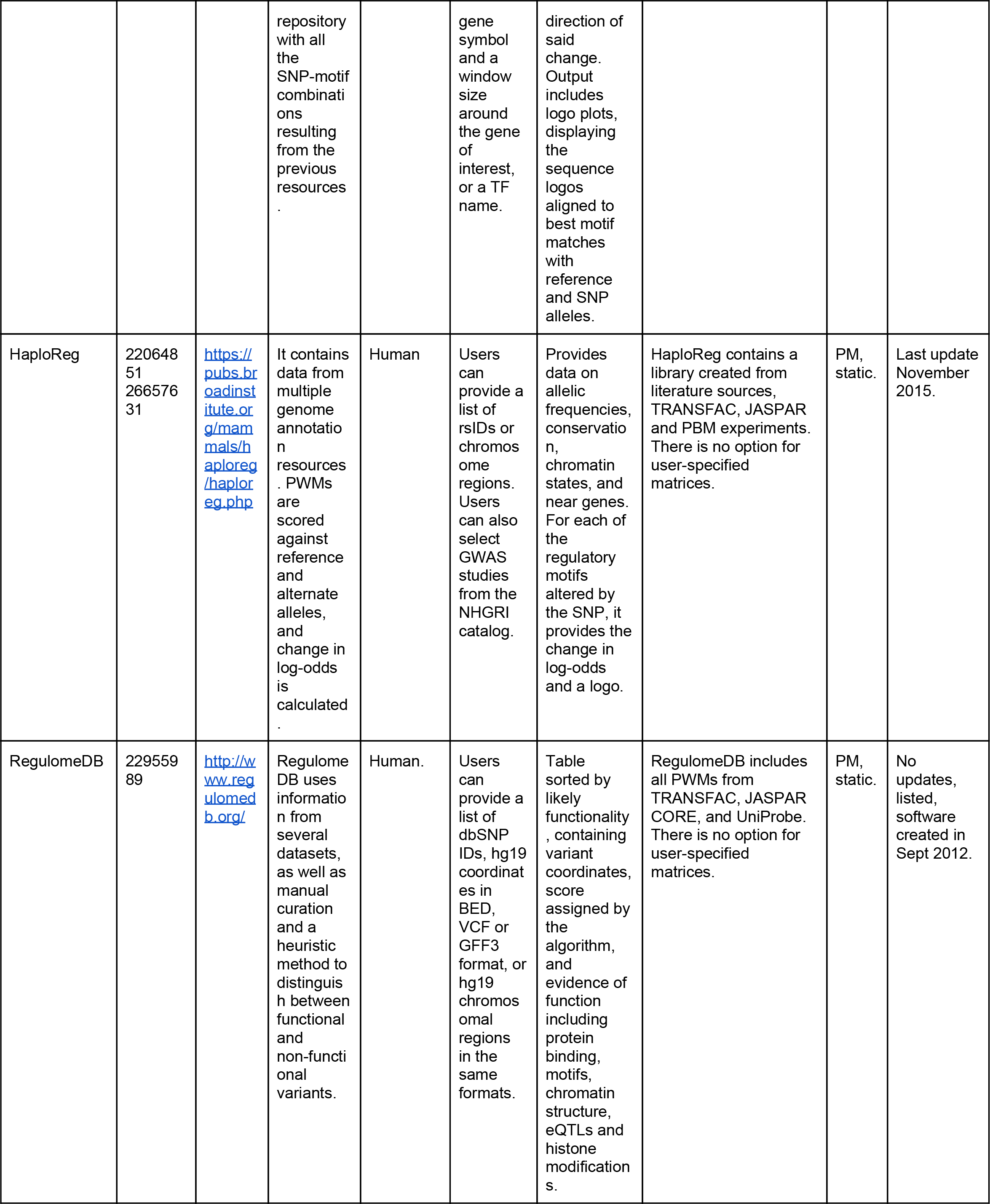
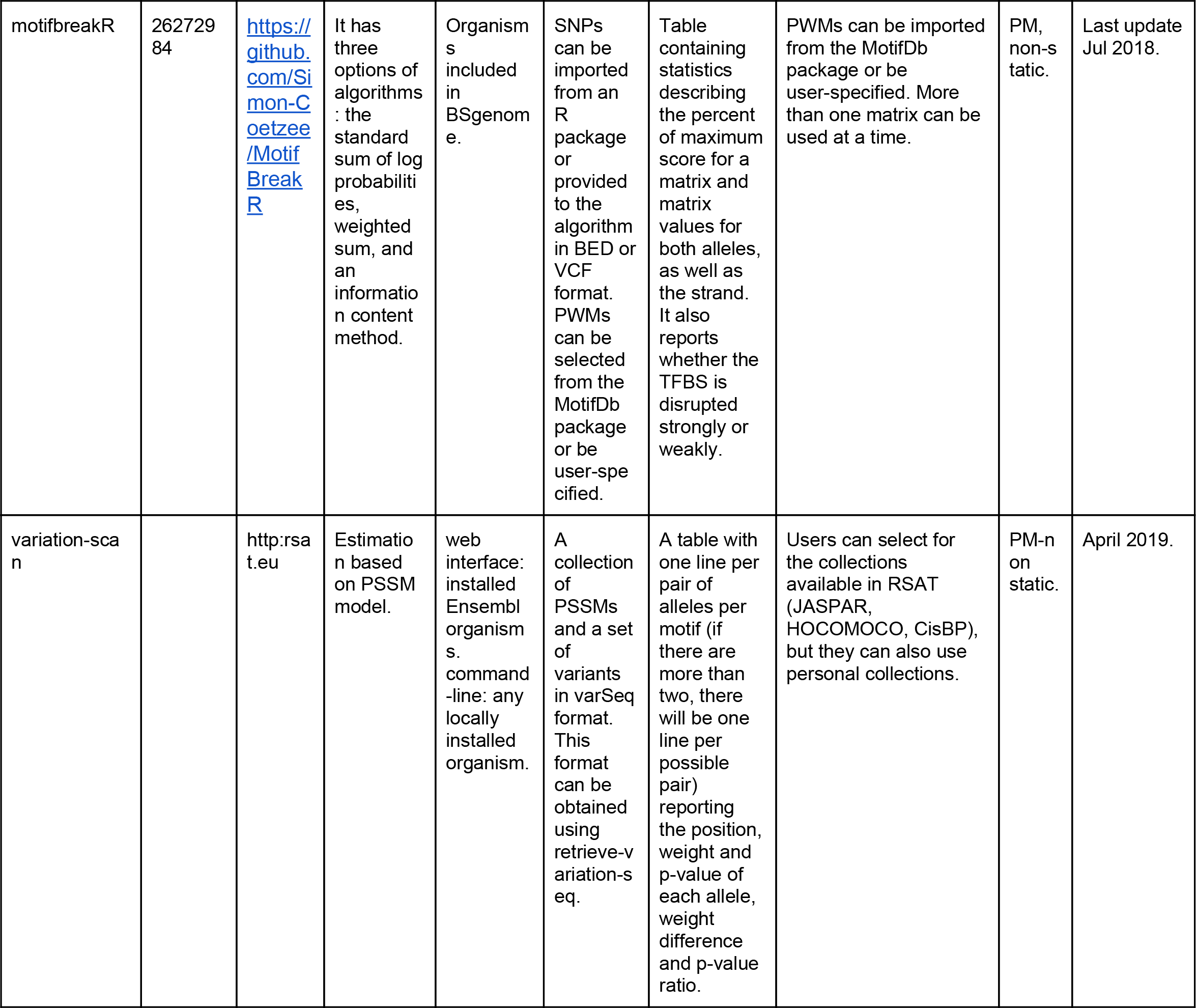
Tools similar to variation-scan with available implementation. PM stands for Pattern Matching, ML stands for Machine Learning

DeepSea [6] integrates functional genomic data from ChIP-seq, DNAseq, RNA-seq and other functional genomic high-throughput data to assess the potential damage of variants across the human genome. Precalculated results for annotated variants can be accessed on their website.

Both tools can be trained on other organisms, provided that functional genomic data are available. The main limitation of these resources is the required expertise in bioinformatics and/or computational resources for users to analyse their own data sets.

Another set of tools identify potential regulatory effects of a variant by comparing the measured affinity of a TF to the different possibles alleles. Our tool, named *variation-scan*, falls within this category.

All tools in the Pattern Matching category, (labeled PM in Table1) use Position-Specific Scoring Matrices (PSSMs) to evaluate the affinity of a TF to a given sequence with an allele. Major differences between these tools can be found in (i) their availability: web pages [9], command line [14] or both [15]); (ii) flexibility for the user to input their own data [16]; (iii) usability (possibility to use several variant formats [7]); (iv) results representation: figures and/or tables [10]; (v) available organisms: only human [17], or other organisms [16]); and (vi) the possibility to calculate results on-the-fly [12] or access pre-calculated ones [9]. One of the major challenges, aside from the development itself, is the maintenance over time and tool updating.

### Var-tools

In this context, we have developed *Var-tools* to address the main limitations identified in existing programs (Table 1). *Var-tools* are composed of four programs that enable (i) retrieval of information of Ensembl annotated variants when available for a given genome in RSAT (*variation-info*), (ii) conversions between variant file formats (*convert-variations*), (iii) retrieval of the sequences surrounding variants (*retrieve-variation-seq*), and (iv) scanning of different alleles of a variant with one or several motifs, comparing the scores and p-values in order to identify affected TFBSs (*variation-scan*).

In summary, RSAT *Var-tools* provide an accessible resource for experienced and non-expert users to analyze regulatory variants in a web interface for fifteen organisms (five metazoa (http://metazoa.rsat.eu) and ten plants (http://plants.rsat.eu), with flexibility to upload personal variant and PSSM collections. We describe here *Var-tools* methodology, along with four case studies demonstrating the flexibility of the tools, enabling the analysis of data sets from different origin (Ensembl variants, Genome-Wide Association Study (GWAS) data, ChIP-seq regions, etc.), complexity, and organisms.

## Methods

### Var-Tools: From variants to identification of regulatory effects

Var-tools consist in a subset of four tools within RSAT devoted to the identification of genetic variants putatively affecting TF binding.

1. *variation-info*: this tool relies on the Ensembl genetic variation information [18] annotated and installed on the corresponding server for each particular genome (*i*.*e*. Human variants are installed in the Metazoa server). It can take two different inputs variant rsID or 2) genomic loci in bed format. This tool will retrieve the information of the variants matching the IDs or the information of the variants located in the genomic loci. Variants installed in RSAT servers have been processed to remove variants with incomplete annotations (no alleles) or ambiguous coordinates (alleles coordinates don’t match). When users have their own variants collections, they can skip this tool and use directly *convert-variations*.
2. *convert-variations*: enables the interconversion of variant files format such as VCF, GVF and varBed. varBed is an internal format of RSAT that facilitates the retrieval of the sequence surrounding the variant (Supplementary Figure 1A).
3. *retrieve-variation-seq*: retrieves the sequence surrounding the variant, and produces one sequence for each allele (Supplementary Figure 1B). The tool can take as input a varBed file (see *convert-variations*), and for organisms with Ensembl annotated variants, it can take a list of IDs or a bed file listing genomic loci. The output is provided in a format named varSeq, with each row giving one allele with its surrounding sequence. Each variant has a specific internal ID to accommodate several variants with various alleles in the same file.
4. *variation-scan*: performs the scanning of alleles with a PSSM and compares the scores and p-values between alleles to assess the putative effect on TF binding (see details below) (Supplementary Figure 2). It requires as input a varSeq file (see *retrieve-variation-seq*), a motif or collection of motifs (over twenty supported file formats), and a background model (for methodological details on background model, refer to [19] Box n°3). Different background models are readily available through the web interface. However, depending on the biological question and related potential biases, we recommend the creation of a dedicated background model, which can be done using the RSAT tool *create-background*, also available via the RSAT web interface.

### Haplotype Processing

Genetic variants can be detected using high-throughput techniques. This has enabled the identification of millions of variants in the HapMap [20] and 1000 genomes projects [21]. However, the information on the variants alone is less useful than knowing which groups of alleles are co-located on the same chromosome (haplotype). The process of identifying the variants that belong to each chromosome is known as phasing. Including haplotype phasing information facilitates the identification of relations between variants [22].

VCF files can include haplotype phasing information. The tool *convert-variations* identifies and retrieves the phasing information of the variants, while the tool *retrieve-variation-seq* builts the corresponding haplotype with all the SNPs that lay within a defined window (default: 30 bp).

### Computing binding specificity of a transcription factor to a DNA sequence

*variation-scan* relies on using PSSMs to assess the binding specificity of a TF to a DNA sequence with different alleles in a given position. The first step of *variation-scan (i*.*e*., scanning of the sequences with a given PSSM) is internally delegated to the RSAT tool *matrix-scan*. The scoring scheme and p-value calculation are described in detail in [19], Box n°1 and Box n°2, respectively. In brief:

PSSM are used to assess the binding specificity of a TF, this affinity is calculated as a weight score (*Ws*). The *Ws* of a site in *variation-scan* is calculated using [23]:

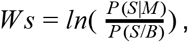

where S is a sequence segment of the same length of M, M is the PSSM, and B is the background model. Hence, P(S|M) is the probability of the sequence given the PSSM and P(S|B) is the probability of the sequence given the background model.

Moreover, it is possible to calculate the p-value of a given score as:

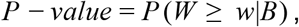

where the P-value is calculated as the probability of observing a score of at least *W* given a background model *wlB*.

When a sequence is longer than the PSSM, the PSSM is shifted base by base until the full sequence has been scored. This scanning step is performed on the sequences of all reported alleles, so that each allele is compared with all the positions of a with a given motif.

### Assessment of allele effect on transcription factor binding

In the second step (i.e., evaluating the impact of SNPs), *variation-scan* compares the obtained Ws (Ws difference= Ws_Allele1 - Ws_Alelle2) and the *P-value* (*P-value* ratio= P-value_Allele1 / P-value_Allele2) of each of the alleles, position by position throughout the scanning window. To evaluate indels, *variation-scan* compares the highest Ws and its corresponding *P-value* for each sequence of the reported alleles. When more than two alleles of a variant are reported, all alleles are compared to all alleles in a pairwise manner.

### variation-scan performance test

#### Computing efficiency

The tools *variation-info* and *convert-variation* are programed in Perl, while *retrieve-variation-seq* and *variation-scan* are coded in C, to enable the analysis of large numbers of variants from eukaryotic genomes in a reasonable time. To further improve performance, we reduced the data transfer from the hard drive to memory.

*variation-scan* performance was assessed by randomly selecting a variant from the 1000 genomes project [21] and a motif from the RSAT non-redundant motifs collection [24]. The randomly selected variant was used to create sets with different numbers of replicates, ranging from one thousand to nine millions, to estimate the relation between running time and the amount of evaluated variants.

#### Dataset: experimentally-determined regulatory variants in red blood cells

The regulatory activity of 2756 red blood cell variants has been systematically measured using Massively Parallel Reporter Assays (MPRA) [25]. MPRA is a high-throughput assay in which a library of putative regulatory elements, each followed by a unique barcode, is inserted into a plasmid, then transfected into a cell, and quantified through the abundance of barcodes. These variants are known to be in strong linkage disequilibrium (LD) with 75 variants associated with common traits of this cell type. Three sliding windows per variant (left, right, and center) were synthesized, barcoded and used for to study the effect of slight changes in their genomic context. For each sequence mRNA/DNA ratio was computed to obtain a quantitative evaluation of the regulatory effect a sequence variant.

#### Evaluation of variation-scan

The variant dataset was used as input for *variation-scan*; the variants assessed in the red blood cell assay were annotated with the Ensembl GRCh37 human genome release, and given as input to *convert-variations* followed by *retrieve-variation-seq*. Since three sliding windows were used for each variant in the MPRA, the corresponding windows were merged before computing a background model using the *create-background-model* tool.

According to the original study [25], binding sites for the following TFs were enriched in the sequences of interest: GATA1, KLF1, DHS, TAL1, ETS, FLI1 and AP-1. Therefore, a total of 48 PSSMs annotated as related to these TFs were retrieved from the non-redundant RSAT motif collection [24], and given as input to *variation-scan*.

A negative control set of motifs was created using the RSAT tool *permute-matrix* [4]; five pemuted motifs were created for each of the 48 motifs, generating a collection of 240 control motifs.

For a variant to be reported in *variation-scan* as positive, we requested that at least one of the allele sequences was evaluated as a binding sites with a p-value of at most 10^−4^(using the parameter -uth pval 1e-4 in the command line), and that the p-value ratio was greater or equal to ten (a change of one order of magnitude between the best and the worst allele p-values) (-lth pval_ratio 10).

### Case studies

#### Case study 1: Identification of regulatory variants in the “Platinum” Genomes haplotypes

The set of high-confidence variants from the two CEU (Northern Europeans from Utah) human Platinum Genomes NA12877 and NA12878 [26] were downloaded through the Amazon Web Service (AWS) Command Line Interface from the Illumina Platinum Genomes AWS S3 bucket (https://github.com/Illumina/PlatinumGenomes). The downloaded VCF files contained phasing information of each CEU individual haplotype configuration. The genome version used was GRCh37.

We selected SNPs intersecting with the annotated DNAse-seq clustered peaks V3 from the ENCODE project [27]. The VCF file with the selected SNPs was processed using *convert-variations* with the option *phased* and then the haplotype sequences were reconstructed with *retrieve-variation-seq*.

For an haplotype SNP set to be reported in *variation-scan*, we requested that at least one of the haplotype sequences was evaluated as a binding site with a p-value of at most 10^−4^(-uth pval 1e-4) and that the p-value ratio between the two haplotypes was greater or equal to 100 (a change of two orders of magnitude between the best and the worst haplotype alleles p-values) (-lth pval_ratio 100). In addition, we require a change of sign between the best and worst score as an additional filter. Finally, single position variants were discarded.

#### Case study 2: prediction of regulatory variants associated to the susceptibility to Mycobacterium tuberculosis

We collected SNPs associated to the phenotypic trait “*susceptibility to Mycobacterium tuberculosis infection measurement*” (disease ID EFO_0008407) from the latest version (1.0.2) of the GWAS catalog [2] (https://www.ebi.ac.uk/gwas/). This query returned one study [28] with 67 distinct variants, of which 48 had a valid reference SNP identifier (rsID) and could be further used (denoted hereafter as disease-associated SNPs, or DA-SNPs). To predict the TF binding sites putatively affected by these selected SNPs, we designed an approach combining Var-tools with different external resources. We further collected from Ensembl REST interface (http://rest.ensembl.org/) 564 SNPs in linkage disequilibrium (LD-SNPs) in the European population [17], with a threshold on the regression coefficient (r^2^ ≥ 0.8) and a maximal distance of 200 bp.

Annotations (chromosomal location, type of genomic region) of the resulting 612 SNPs (48 DA + 564 LD) were collected from Ensembl BioMart [29,30]. We then selected SNPs in non-coding regions, resulting in a set of 572 SNPs of interest (SOIs) for the detection of regulatory variants. Using SNPs in LD we determined LD-Block regions, we then filtered the these regions overlapping them with ChIP-seq peaks, and calculated the enrichment of TF binding peaks using ReMap [31], we then calculated enrichment for disease annotations using R XGR package[32].

Finally, we used *retrieve-variation-seq* to retrieve the sequence variants around each SOI, and predicted the impact of the variation on TF binding for each motif of the JASPAR non-redundant RSAT motif collection [24] using *variation-scan*, with a threshold of 1e-4 on the p-value and 100 on the p-value ratio.

#### Case study 3: Assessment of the regulatory effect of GWAS reported variants in promoters with enhancer function

The STARR-seq assay [33] is in its principle similar to MPRA, and helps identify self-transcribing active regulatory regions that have *enhancer* potential. Using this approach, Dao *et al* (2017) [34] analysed the *enhancer* potential of annotated RefSeq promoters [35]. In the two cell lines K562 and HELA, they identified 632 and 493 promoters with enhancer function (ePromoters), respectively. Moreover, the authors identified enrichment of eQTL variants reported by GTEx [36].

To identify ePromoters variants that could be affecting TF binding, we retrieved the GWAS catalog version v1.0 (downloaded on 7/01/19) [2] and overlapped it with the ePromoter coordinates reported in [34], using bedtools overlap (version 2.26.0) [37]. PSSMs representing TFs enriched in ePromoters were also obtained from [34], corresponding to SMRC1, JUN, FOS, ATF:MAF:NEF2, YY1, ETS family, Creb and USF1/2.

Using the selected GWAS variants that fall within ePromoters and the motifs for enriched TFs in those regions, we applied *variation-scan* to assess the potential regulatory effect of these variants. *variation-scan* was run with the parameters -lth w_diff 1 -lth pval_ratio 10, with a background model built using *create-background* with all RefSeq promoter sequences. In order to filtrate variants with the highest putative regulatory disruption, we further selected variants that showed a change of sign in the weight score between alleles.

#### Case study 4: Identification of regulatory variants affecting VRN1 binding in barley

The latest version of the *Hordeum vulgare* (barley) reference genome [38] and a panel of mapped genetic variants were imported from Ensembl Genomes release 42 [39] and installed in the RSAT Plants server (http://plants.rsat.eu). We obtained experimentally determined binding sites (ChIP-seq) for VRN1 from [40]. Since these peaks were originally positioned within contigs of the 2012 genome assembly [41], they had to be matched to the corresponding regions of the current assembly with BLAST+ v2.9.0 (blastn) local alignments against the repeat-masked genome sequence (perfect matches) [42]. Using bedtools overlap (version 2.26.0) [37], we selected variants falling within the VRN1 reported binding peaks. The selected variants in VCF format were then processed using *convert-variations* and *retrieve-variation-seq* to obtain the sequences with the alternative alleles.

The VRN1 DNA motif used to scan the variants was obtained from the footprintDB plant collection [43] version: 2018-06 (http://floresta.eead.csic.es/footprintdb/index.php?motif=AY750993:VRN1:EEADannot). *variation-scan* was used with a pre-computed background Markov model (order 1) for barley to assess the effect of variants in TF binding, with the following parameters: -lth score 1 -lth w_diff 1 -lth pval_ratio 10 -uth pval 1e-3.

## Results

The *Var-tools* provide companion programs enabling the retrieval of variants (*variation-info)*, their surrounding sequences (*retrieve-variation-seq*) and interconversion between file formats (*convert-variation*). The main predictive program is *variation-scan*, which can be used with any set of variants provided by the user (in vcf or gvf formats) or annotated in Ensembl (from a list of rsIDs or a bed file to identify overlapping variants in genome coordinates), with any set of motifs selected from the collections available in RSAT, or provided by the user.

### variation-scan accurately assesses the effect of experimentally validated regulatory variants

The original version of *variation-scan [44]* required approximately five hours to assess the allele effect of nine millions variants. The novel version [4] significantly reduces the processing time to about one hour (Supplementary Figure 3).

To evaluate the performance of *variation-scan*, we used an experimentally validated regulatory variant set obtained from a Massively Parallel Reporter Assays (MPRA) experiment [25]. For all of the assessed allele pairs, we compared the obtained weight score difference from *variation-scan*, to the mRNA/DNA ratio of the MPRA (see methods). As shown in Figure 2A, we are able to recover 96.8% of the experimentally validated variants with *variation-scan*. Using the variants reported as positive in the MPRA data set, we were able to observe correlation between the weight difference and the MPRA mRNA/DNA ratio in positive variants, showing that *variation-scan* gives accurate measurements of the impact of regulatory variants (Figure 2 B). With the proposed thresholds, we can confidently reject 31.5% of MPRA negative sequences, while this could be improved using more restrictive parameter, this would imply a reduction in true positives. Moreover, as any hightrouput assay, MPRA has its limitations [45] and sequencing biases could increase the number of false negatives.

**Figure 1.**
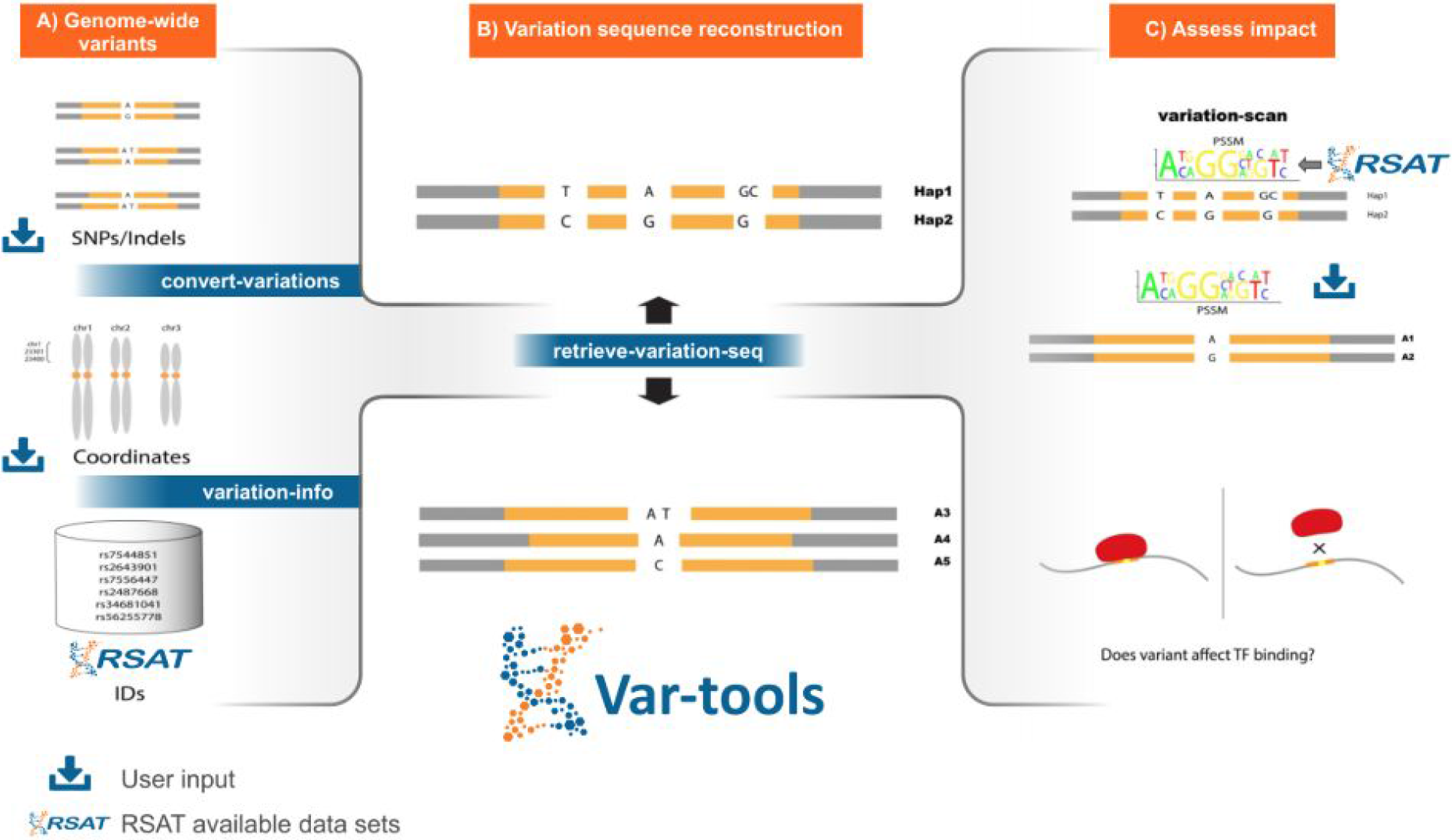
Schematic representation of Var-Tools: This set of tools included in RSAT is focuses on assessing the impact of different allelelic variants on transcription factor binding sites. A) *convert-variations* allows users to input their own variants and convert them to other formats (gff and varBed, the latter is the format used in the next step), while *variation-info* retrieves the annotated information of Ensembl variants installed in RSAT servers. B) The tool *retrieve-variation-seq* retrieves the surrounding sequence of variants (including possible haplotypes) and generates a text file with one line per allele and per variant or haplotype (varSeq format). C) Users can input their variants in varSeq format and a collection of motifs (direct input by the user or selected from RSAT available collections) to *variation-scan*; the tool will scan with all motifs and compare in a pairwise manner each allele of a variant or haplotype.

**Figure 2:**
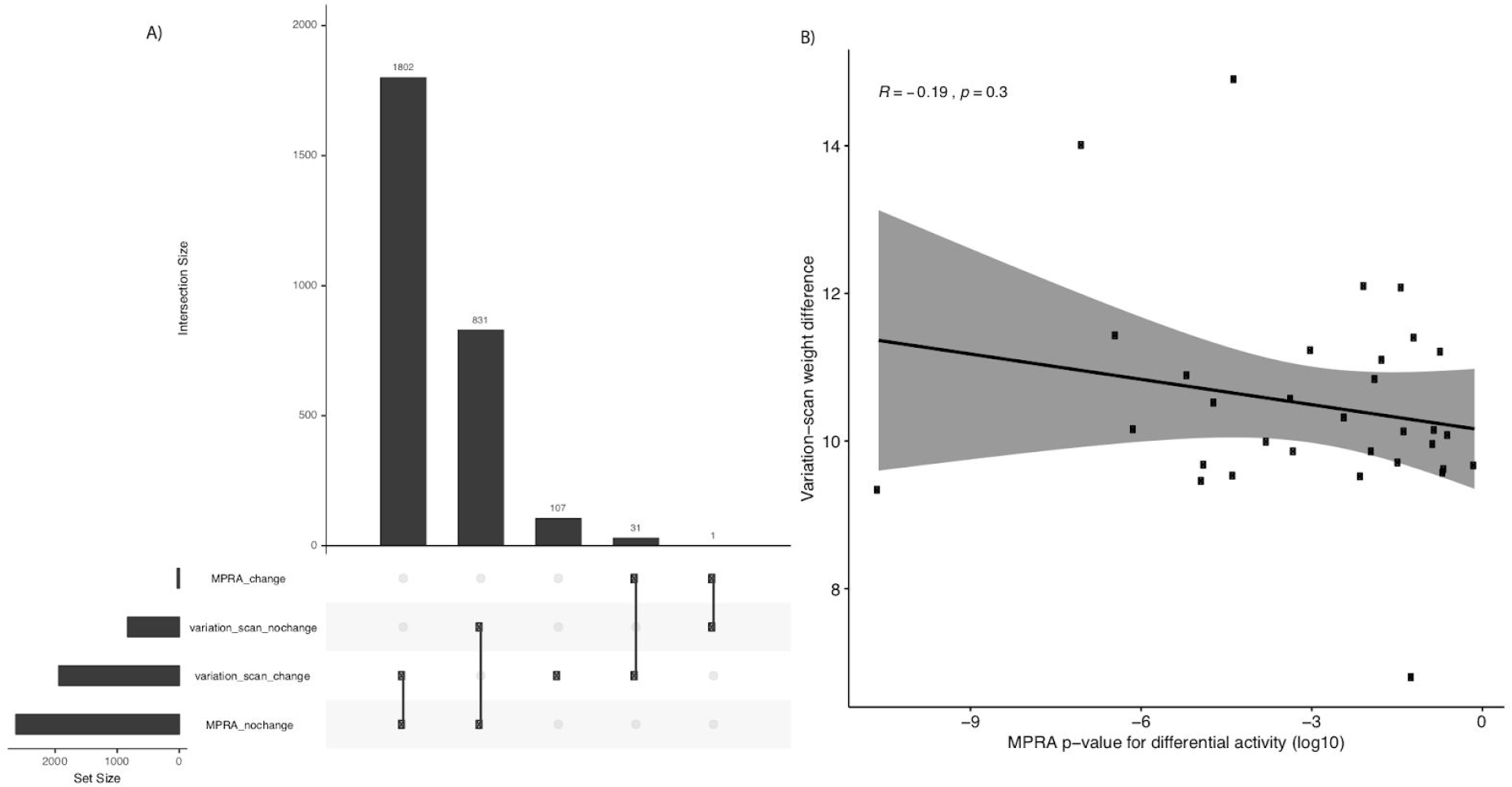
Identification of experimentally validated regulatory variants using *variation-scan*. A) Histogram showing the number of MPRA and *variation-scan* evaluated variants. Variants were classified in both cases as “change” or “no change”, depending on whether they showed an effect caused by the variant on the MPRA assay or in the *variation-scan* evaluation. B) Correlation of the MPRA p-value of the mRNA/DNA ratio of positive variants and the *variation-scan* weight difference for the MPRA variants with significant change.

We performed a negative control, consisting of 240 permuted matrices (five permuted versions of the 48 motifs). With this collection, it was still possible to recover a group of variants, but it only represented 37.8% of the MPRA positive variants (Supplementary Figure 4).

### var-tools case studies

To illustrate the diverse applications of *var-tools* to tackle various biological questions, we designed three different case studies:

1. Impact of regulatory variants in the same haplotype on TF binding sites.
2. Identification of the regulatory potential of variants reported in GWAS.
3. Assessment of the regulatory potential of GWAS variants within experimentally determined regulatory regions.
4. Determination of regulatory variants within TF binding regions identified using ChIP-seq [40].

#### 1 Genome-wide haplotype variant information can be used to identify sets of regulatory variants affecting the same TFBS

The lowering costs in sequencing have made it possible to obtain whole genomes of more individuals, opening the possibility of knowing, not only the variants of a genome, but also the haplotypes, and determining which variants are passed linked within the same chromosome. This enables now the possibility of assessing the regulatory effect of sets of variants within the same haplotype in a given TF binding site.

Using the high-confidence SNPs from two “Platinum” Genomes [26], we determined haplotype variants that are likely to affect one TF binding site. We selected variants 30bps apart, located in open chromatin, to be analysed with *variation-scan* using the non redundant motif collection at RSAT [24]. We detected 7406 haplotype sites with more than one SNP with a probable effect in binding of 361 TFs. Overall the number of heterozygous variants within an haplotype increases the measured weight difference, this is expected as more changes in the binding sites are more likely to change TF affinity (Figure 3A).

**Figure 3:**
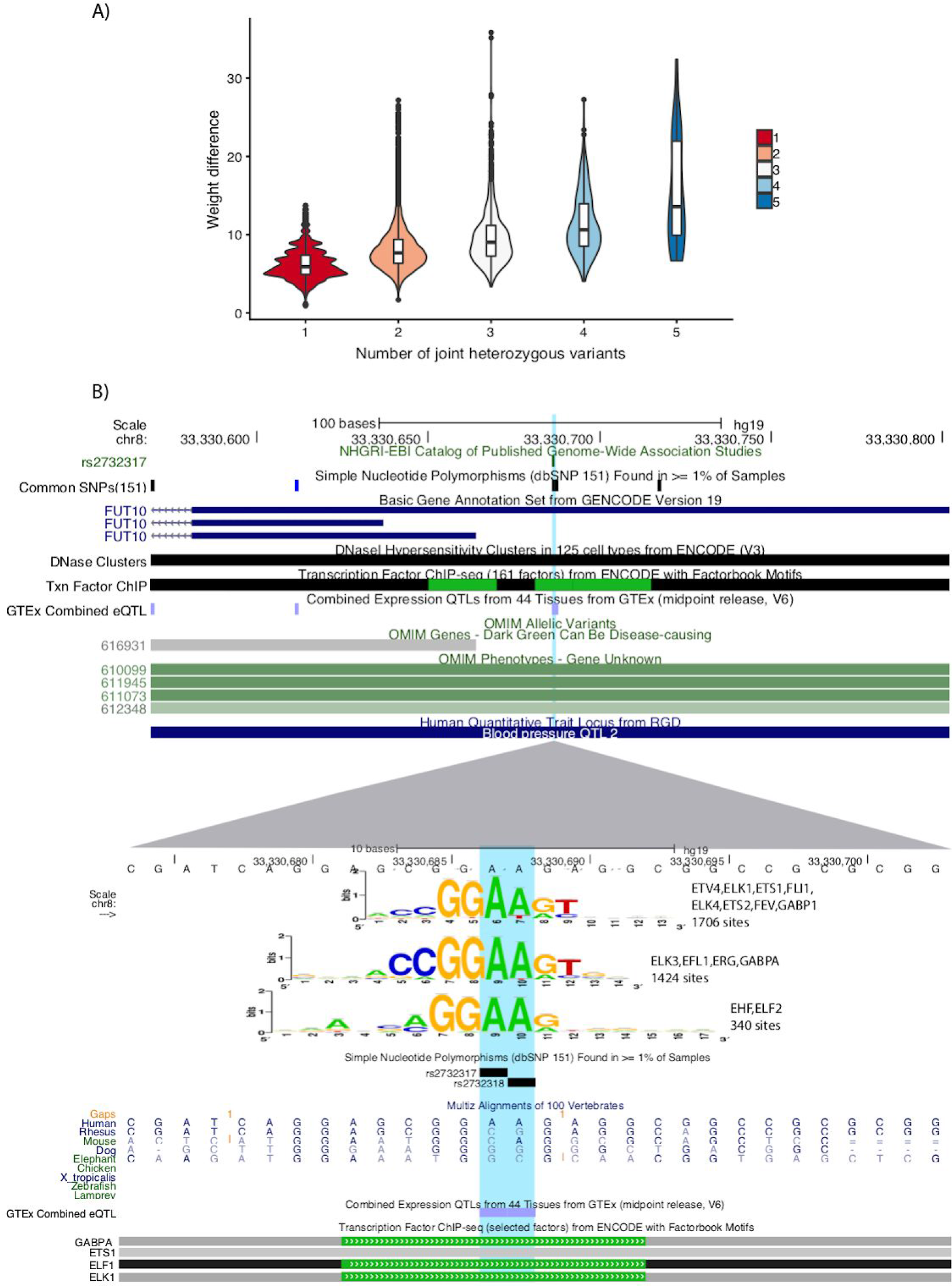
Haplotype analysis in high-quality human genomes. A) The number of heterozygous variants (x axis) within the same putative binding site will tend to have a greater impact on the TF binding probability when comparing haplotypes (weight difference, y axis). B) UCSC browser [48] screen shot, showing the locus of two SNPs that compose an heterozygous haplotype in one of the CEU individual, figure show the reference genome haplotype. The variants are located in the FUT10 promoter (top), *variation-scan* predicts an effect in three motifs that represent binding sites for GABPA, ETS1, ELF1 and ELK1, factors that have been proven to have binding sites in this region by the ENCODE project. The variant rs2732317 has been associated with effects in gene expression.

One of these haplotypes is composed of the minor alleles of two SNPs (rs2732317 and rs2732318), where we observed a potential regulatory effect likely affecting binding of three motifs: EHF/ELF2, ETV4/ELK1/ETS1/FLI1/ELK4/ETS2/FEV/GABP1, and ELK3/ELF1/ERG/GABPA. This finding is consistent with TF binding reported by the ENCODE consortium in this region (Figure 3B).

#### 2 Genetic variants associated to Mycobacterium tuberculosis infection show potential regulatory effects

The second case study illustrates a knowledge-free use of *Var-tools* to identify regulatory variants from GWAS studies for a user-specified disease, without prior indication about the potentially involved transcription factors or binding motifs. The approach is based on the prediction of regulatory variants with RSAT *Var-tools*, narrowed down by selecting the regulatory SNPS that overlap ChIP-seq peaks in ReMap[31], in order to identify convergent indications for a potential impact of the variants on the binding of a TF.

The 572 SNPs of interest (SOIs, see methods) show a significant enrichment for diseases related to respiratory functions (lung carcinoma, respiratory neoplasm, squamous cell carcinoma, lung disease) as well as for schizophrenia (Figure 4), confirming the relevance of the collection of SNPs.

**Figure 4.**
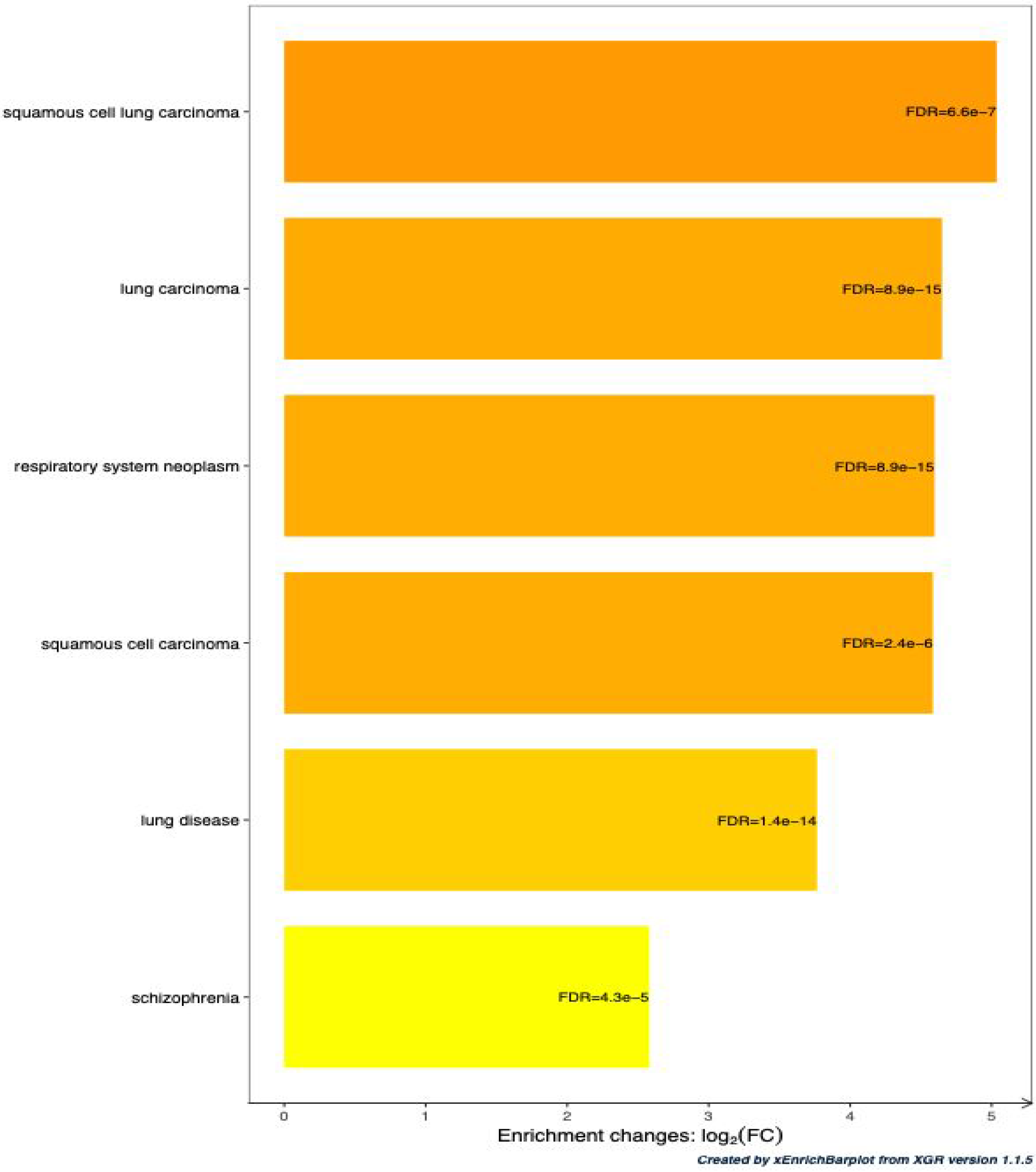
Enrichment of the set of SOIs for diseases. Comparison of the significant phenotype ontology terms.

The scanning of the SOIs with the 1,404 matrices of Jaspar core non-redundant Vertebrate collection predicted 363 modification of TF binding, covering 145 distinct SNPs and 181 distinct motifs. There are 2,882 overlaps between the 35,393,642 ChIP-seq peaks of the ReMap catalog and the 572 SOIs, but only two of them show a match between the TF of the ChIP-seq peak and the TF associated to the *variation-scan* motif: CEBPB for reference SNP rs3131071 and ELF1 for rs3132397. Noticeably, CEBPB has been reported as main regulator for genes differentially expressed between tuberculosis patients and control cases [46]. In summary, the convergence of ChIP-seq and motif scanning thus enabled to identify 2 priority candidates among the relatively large list of 145 candidate regulatory SNPs. The same approach can be applied to other association studies in order to predict regulatory variations potentially involved in user-specified diseases.

#### 3 Assessment of the regulatory effect of GWAS reported variants in promoters with enhancer function

ePromoters are regulatory regions with dual functions: as promoters, they regulate the gene downstream, but they also show *enhancer* potential when tested for self transcription [34], indicating that they could act as an *enhancer* for other genes. ePromoters have been described to be enriched for eQTLs, which means their function could be affected by genetic variants. We thus set out to investigate if GWAS reported variants could be affecting TF binding in ePromoters.

We identified five and twelve GWAS reported variants falling within the reported coordinates of ePromoters corresponding to the human cell lines HELA and K562, respectively. Using *variation-scan* with the motifs of TFs reported as enriched in ePromoters, we were able to identify effect of two variants (rs3771180, rs3822259) on the binding of two TF motifs in HELA (MAF::NFE2, FLI1/FEV/ETS2/ELK4/ELK4/GABP1/Gabpa), and two variants (rs147997200, rs62229372) affecting three binding motifs in K562 (Creb3l2, Atf3/MAFG::NFE2L1/MAFG/NF2L1, FOS::JUN).

Noteworthy, in HELA ePromoters, we found the SNP rs3771180, which is also described as an eQTL in the whole blood dataset by the GTEx project. This variant has been associated to asthma and is upstream the Interleukin Receptor 1 gene, which is consistent with ePromoters being related to inflammatory response in HELA [34].

#### 4 Population genetic variations in barley can potentially affect VRN1 binding

Variation in traits in crops can be due to changes in TF binding that likely affect gene regulation. As proof of concept that *var-tools* can be used for this purpose, we set out to identify reported variants in *Hordeum vulgare* (barley) that can putatively affect binding of VRN1, a TF directly linked with vernalization response. We focused this analysis on VRN1 ChIP-seq reported regions [40], selecting only variants that overlapped them (n=1604).

Using the VRN1 motif annotated in footprintDB and the barley variants annotated in Ensembl Plants, we identified a total of thirteen variants likely to affect VRN1 binding. Of these, two are proximal to genes MLOC_73196 and MLOC_79452, which are among a set of 38 genes observed to change their expression level upon VRN1 binding in RNA-seq experiments [40].

## Discussion

The lowering costs of sequencing technologies has facilitated the identification of genetic variants associated with traits and diseases, in humans and other species. For this reason, the identification of variants affecting TF binding sites has become mainstream, calling for efficient computational approaches to analyse large sets of variants.

One issue arising from the analysis of big data sets is number of false positives. It has been shown that taking advantage of specific biological insights (*e.g*. identification of relevant TFs, reduction of genomic regions using functional genomic information) can significantly improve results, reducing the number of false positives [47,48]. In this respect, the evaluation set and case studies 3 and 4 focused on the analysis on TFs known to bind on the regions of interest, which enabled us to consistently evaluate the performance of the tool in the evaluation set, and further helped us to identify biologically relevant regulatory variants, affecting ePromoters function in case study 3, and VRN1 binding in barley in case study 4. While for case study 2, the usage of ChIP-seq information enabled the identification of potentially relevant variants related to tuberculosis.

Moreover, current genotype information facilitates the characterization of haplotypes, but this requires tools designed to take advantage of this information. The tool *convert-variations* facilitates the usage of this information, enabling the user to analyse the impact of variants within the same chromosome in a 30bp window, potentially affecting the binding site of the same TF. As there was no previous reports of this cases, we decided to use the complete collection of motifs. Nevertheless, by requiring more than one SNP affecting one TF binding site, we were able to identify potentially relevant haplotypes with a regulatory effect. In the advent of new genome-wide characterization in population studies, this function will facilitate the integration of phasing information in the search of regulatory variants.

## Conclusions

*Var-tools* enables the prediction of the effect of sequence variants on TF binding. In addition to reasonable computing time, the focus is put on usability and high flexibility for the user: annotated variants can be retrieved from specific genomic loci, as well as from personal collections of variants, motifs (provided as PSSMs) can be chosen from the collections available in RSAT (JASPAR, HOCOMOCO, CisBP, etc.), as well as from users personal set of PSSM. The tools supports various organisms in selected RSAT servers: currently Metazoa, Plants and Teaching. Only the option for variation retrieval from genomic loci in *retrieve-variation-seq* and *variation-info* are functionalities restricted to organisms with annotated variants in Ensembl. In addition to the web interface, *Var-tools* can also be used on the command line to facilitate analysis of personal data sets. *Var-tools* can be used in combination with external databases, as exemplified with the study of GWAS data. Finally, together with the long-lasting RSAT suite, *Var-tools* programs are continuously maintained and updated.

## Acknowledgements

We thank the persons contributing to the maintenance of the RSAT servers, in particular Laboratorio Nacional de Visualización Científica Avanzada (Mexico) specially Luis Alberto Aguilar Bautista and Jair Garcia Sotelo, the ABims platform in Roscoff, France, Pierre Vincens at the ENS, Paris and Aurora Martín Cotaina from EEAD-CSIC for her help on managing the Plants server. We acknowledge Salvatore Spicuglia for useful comments during the development of the tools. We thank Lambert Moyon and Swann Floc’hlay for providing feedback testing *Var-tools*. We thank Mauricio Guzman for styling the figures.

