## Supplementary Figures for "RSAT Var-tools: an accessible and flexible framework to predict the impact of regulatory variants on transcription factor binding"

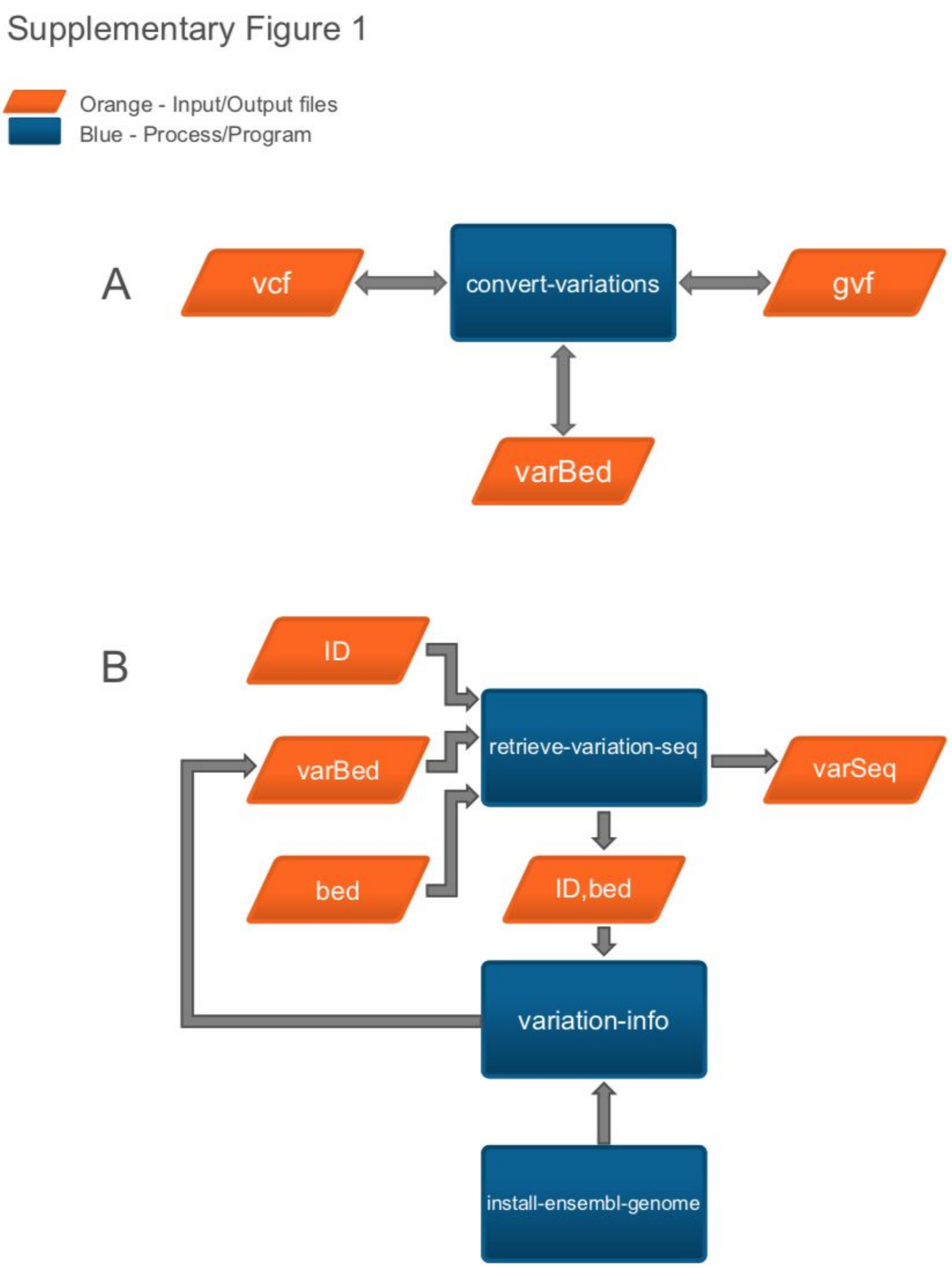


Supplementary Figure 1. **Flowchart describing A) *convert-variations* and B) *retrieve-variation-seq* tools**. RSAT tools are represented by a dark blue rectangle, and inputs and outputs with an orange trapezoid.

Supplementary Figure 2

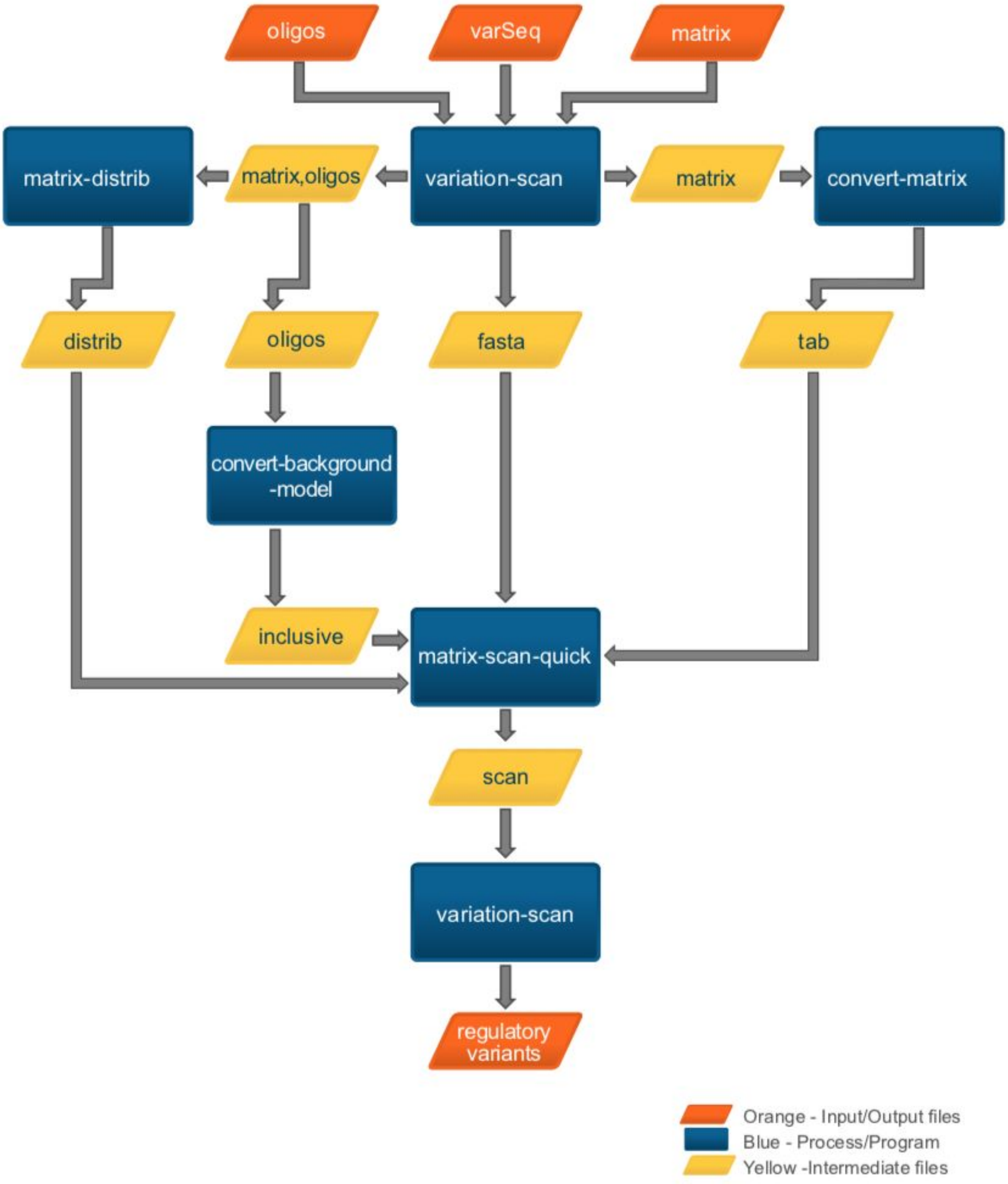


Supplementary Figure 2. **Flowchart describing *variation-scan***. RSAT tools are represented by a dark blue rectangle, and inputs and outputs with an orange trapezoid.

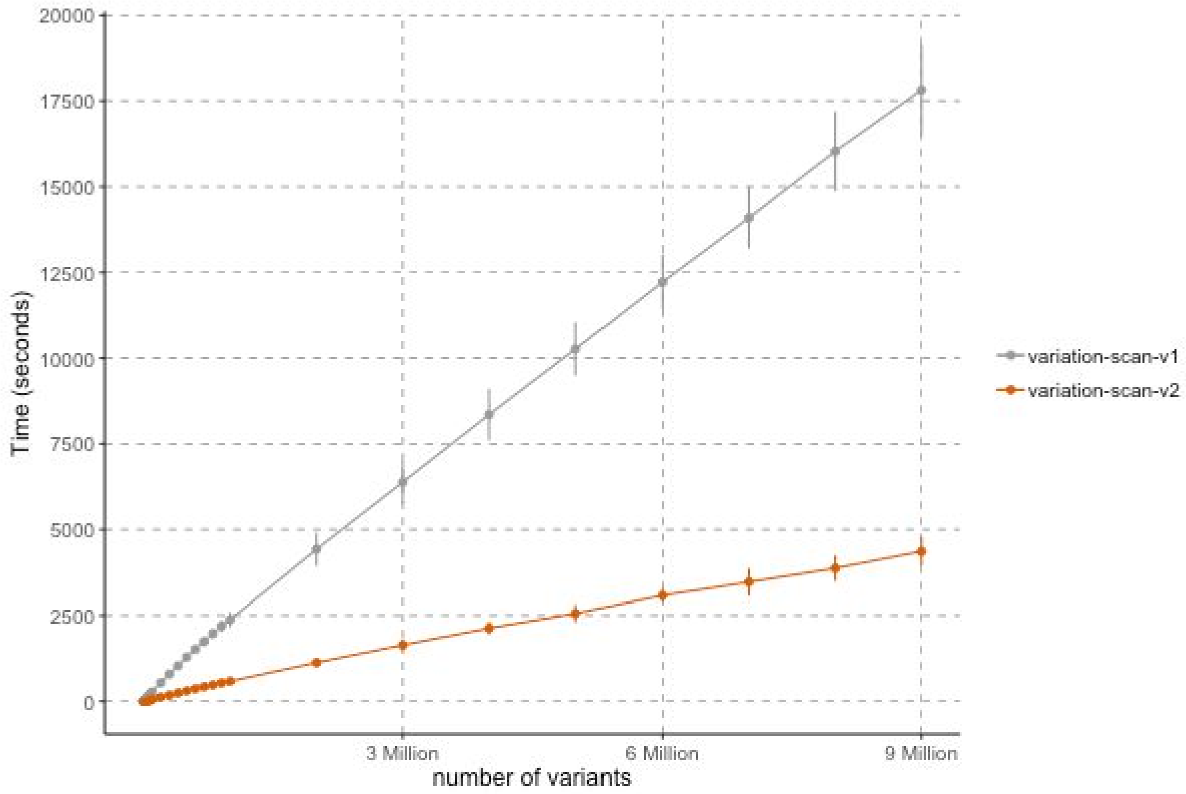


Supplementary Figure 3. **Time efficiency of variation-scan's first and final version**. The abscissa indicates number of variants, the ordinate processing times. The original tool was developed in Perl (grey line). The new version was encoded in C for the RSAT 2018 (Nguyen et al. 2018) release (orange line).

Supplementary Figure 4

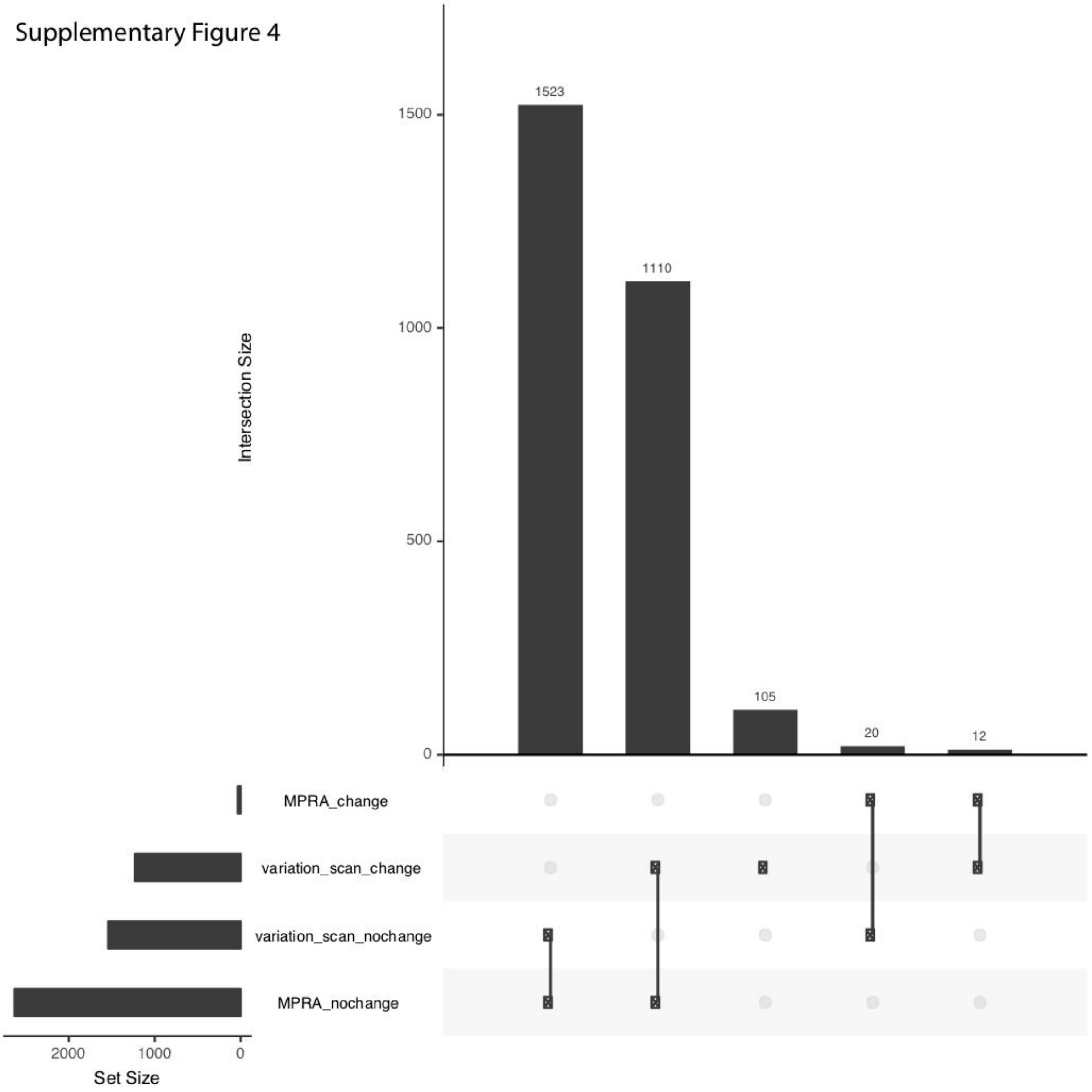


Supplementary Figure 4. variation-scan analyses using motifs with column permutations result in less positive regulatory variants and generate more false positives. Histogram with the interactions between variant group categories MPRA change (positive variants), variation_scan_change (variants evaluated by *variation-scan* as affecting TF binding, see methods for thresholds), variation_scan_nochange (variants evaluated by *variation-scan* has having no effect on TF binding, see methods for thresholds), and MPRA_nochange (negative variants).
